# Genomic regions linked to soft sweeps approximate neutrality when inferring population history from site pattern frequencies

**DOI:** 10.1101/2020.04.21.053629

**Authors:** Nathan S. Harris, Alan R. Rogers

## Abstract

Recent studies have suggested that selection is widespread throughout the genome and largely uncompensated for in inferences of population history. To address this potential issue, we estimated site pattern frequencies for neutral and selection associated areas of the genome. There are notable differences in these frequencies between neutral regions and those affected by selection. However, these differences have relatively small effects when inferring population history.

## 1 INTRODUCTION AND BACKGROUND

In the past year, population geneticists have been debating the extent to which natural selection has shaped the human genome. Evidence suggests that soft sweeps (Harris et al., 2018; Schrider and Kern, 2017) and polygenic adaptation (Daub et al., 2013; Hernandez et al., 2011; Pritchard et al., 2010) are the primary modes of selection in humans. This has led some researchers to suggest that most of the genome is in some way affected by selection either directly or indirectly through linkage with neighboring sites. This led Kern and Hahn (2018) to argue that the original lines of evidence that led to the neutral theory of evolution pioneered by Kimura (1983) do not hold up in the genomic era. Specifically, they believe inferred population histories are skewed because areas of the genome often perceived to be neutral are actually affected by selection. In addition, they suggest genome-wide selection scans will have a high rate of false negatives if using a null distribution built from selected regions of the genome. On the other hand, Harris et al. (2018) and Jensen et al. (2018) suggest that many of these apparent signals of selection are false positives, and others have suggested that selection has been less common but signals of selection are amplified by population history (Torres et al., 2018).

A simple way to the test the idea proposed by Kern and Hahn (2018) is to reconstruct population history using different areas of the genomes. Schrider and Kern (2017) used a machine learning algorithm to assemble a list of regions in the genome inferred to be affected by selection but previously thought to be neutral, and those that are neutral. Here, relative site pattern frequencies (Rogers, 2019) are calculated in each of these subdivisions to measure the potential effects of selection. In what follows, we show that selection skews site pattern frequencies, but has little effect on the estimation of population history in our model.

## 2 RESULTS

### Selection affects population history

A nucleotide site pattern is a particular arrangement of derived and ancestral alleles when a single haploid individual is sampled from each sample population. The total number of distinct site patterns is therefore all combinations in which at least one sample, but not all, carries the derived allele. Three populations are discussed here, given the labels CEU (*X*), JPT (*Y*), and YRI (*Z*). The possible site patterns are *x*, *y*, *z*, *xy*, *xz*, and *yz*, the first three representing the cases in which the derived allele is found in only one population, and the rest representing when the derived allele is found in two populations. Estimations of common ancestry, divergence times, and admixture can be made using the relative frequency of of these site patterns (Rogers, 2019). The difference between the selection-affected site patterns and neutral site patterns was calculated. If neutral and selection-affected genomic regions have similar site pattern frequencies to neutral regions, it will produce similar estimates of population history, and the difference between them should not vary from zero significantly.

Figure 1 shows the difference in relative frequencies of site patterns between affected and neutral regions for biallelic single nucleotide polymorphisms (SNP). Site pattern frequencies were calculated using the *sitepat* program from the *Legofit* package (Rogers, 2019). Confidence in tervals we re ge nerated by using 1,000 bootstrap replicates generated from *sitepat*. Neutral and selection-affected regions differed significantly in the *x*, *y*, a nd *x y* site patterns, with confidence intervals that do not overlap with zero. To investigate the possibility that this pattern is driven primarily by one type of selection, hard sweeps or soft sweeps, affected regions were split accordingly. Hard sweeps differ from other distinctions substantially but the confidence intervals are large, likely due to the relatively small sample size of these regions (Table 1). Soft sweep regions are marginally more similar to neutral regions, but the large difference between affected and neutral regions in the *x*, *y*, and *xy* patterns persists when only soft sweeps are considered.

**Table 1:** Number of sites tabulated in each run of *sitepat*

| Samples | Neutral | Selection Affected | Soft-Sweep Affected | Hard-Sweep Affected |
| --- | --- | --- | --- | --- |
| Humans only | 285,833 | 619,136 | 431,105 | 4,460 |
| Human and Neanderthal | 251,953 | 543,098 | 378,700 | 4,104 |
| Human, Neanderthal, and Denisova | 261,996 | 563,523 | 392,625 | 4,275 |

**Figure 1:**
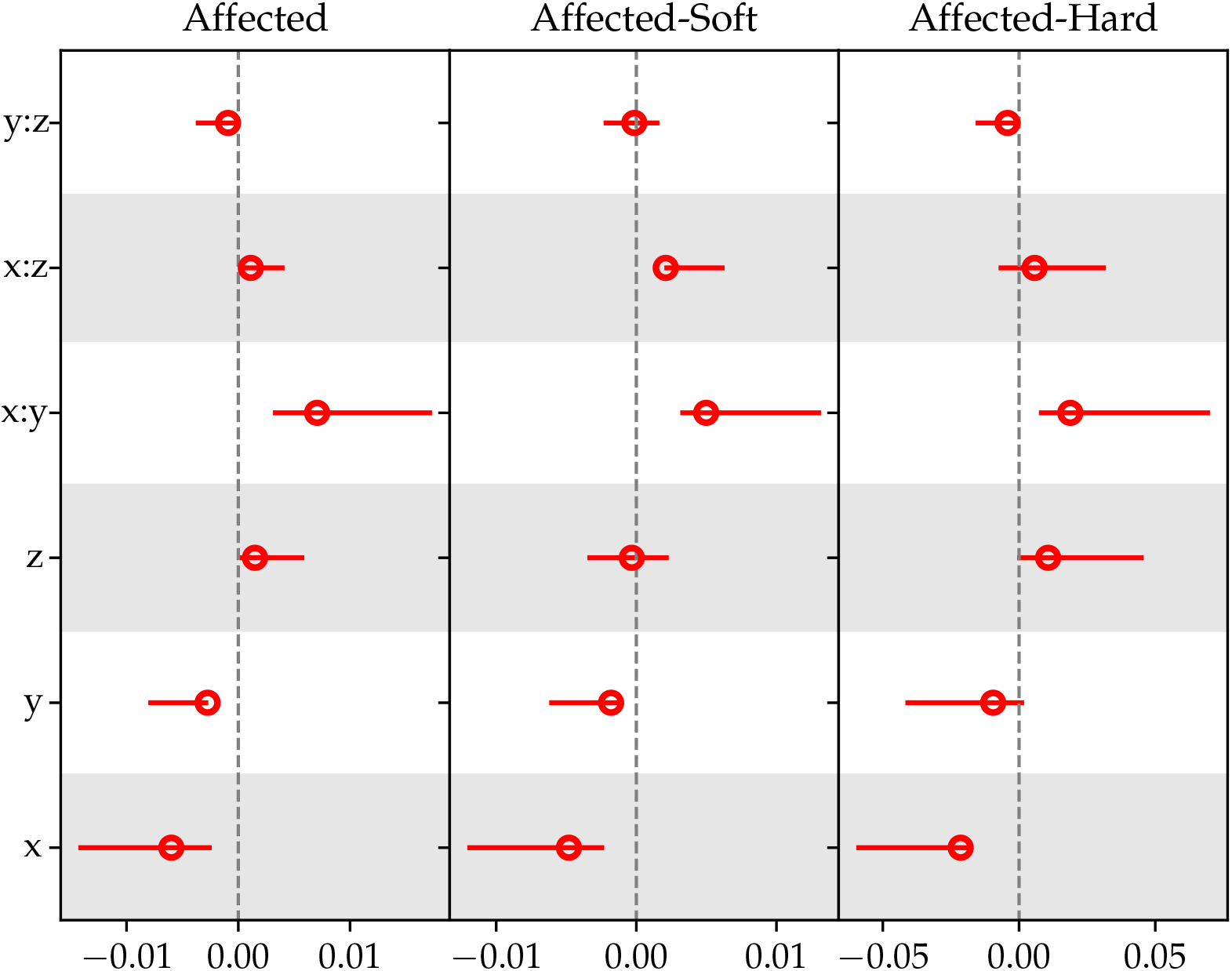
Difference in selection-affected and neutral relative site pattern frequencies. If all patterns rested at zero, the inferred population history would be identical. Each population is given an alphabetical label: *x* (CEU), *y* (JPT), *z* (YRI)

### Inference of Population History

Site pattern frequencies differ between neutral and selected affected regions of the genome. However, these differences are small and it is unclear how large their effect will be when inferring population history. To test the effect these differences have, the site patterns for neutral and soft-sweep affected regions were used to estimate ancestral divergence times and population sizes using *Legofit* (Rogers, 2019). The demographic model used was taken from (Rogers et al., 2019). This model includes admixture events from Neanderthals into Eurasia, ancient humans into Neanderthals, and from superarchaic hominins into Denisovans and the ancestor of Neanderthals and Denisovans. Rogers et al. (2019) found that the exclusion of these admixture events can strongly bias results.

Selection seems to affect the estimation of these parameters, but only to a small extent. Figure 2 shows the percent differences between soft-affected and neutral estimates of divergence times. Confidence intervals were generated by taking differences of individual estimates in each of fifty bootstrap replicates. If selection does not strongly affect quantitative estimates of demographic inference, these differences should rest around zero.

**Figure 2:**
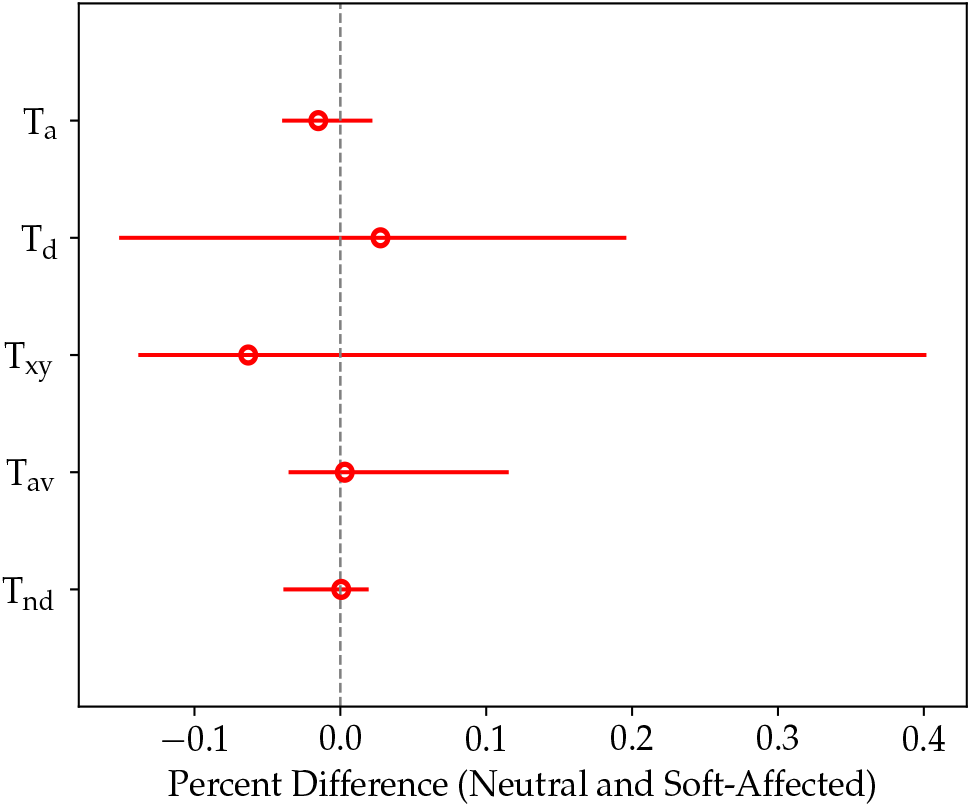
Percent difference between estimates of divergence times for soft-sweep associated regions and neutral regions for a model including Yorubans (*x*), European (CEU) (*y*), Neanderthals (*n*) further split into the Vindija (*v*) and Altai (*a*), Denisovans (*d*), and superarchaic hominins (*s*).

Figure 3 shows the percent differences between soft-affected and neutral estimates of ancestral population sizes. In general, estimates of population size were more likely to be disrupted by the effects of soft selective sweeps. How-ever, these deviations are relatively small. The largest difference in population size without confidence intervals overlapping with zero is approximately one or two percent of approximately 40,000 (Table 2).

**Table 2:** Estimates of divergence times and population sizes for a model including Yorubans (*x*), Europeans (CEU) (*y*), Neanderthals (*n*), Denisovans (*d*), and superarchaic hominins (*s*). “T” indicates divergence time and “2N” is the diploid effective population size.

| Parameter | Neutral | Soft Selection Affected |
| --- | --- | --- |
| $T_{xynds}$ | 67,049.1 | 52,396.2 |
| $T_{nd}$ | 24,906.3 | 24,921.1 |
| $T_{av}$ | 15,570.6 | 15,617.5 |
| $T_{xy}$ | 3,864.6 | 3,628.0 |
| $T_d$ | 1,855.0 | 1,906.8 |
| $T_a$ | 4,958.1 | 4,883.7 |
| $T_v$ | 2,339.7 | 2,315.4 |
| $2N_{av}$ | 31,673.6 | 24,333.9 |
| $2N_n$ | 7,372.7 | 7,414.5 |
| $2N_{nd}$ | 2,111.9 | 2,063.7 |
| $2N_{xy}$ | 50,259.1 | 50,639.3 |
| $2N_{xynd}$ | 42,152.5 | 42,826.4 |
| $2N_s$ | 81,794.2 | 50,639.7 |

**Figure 3:**
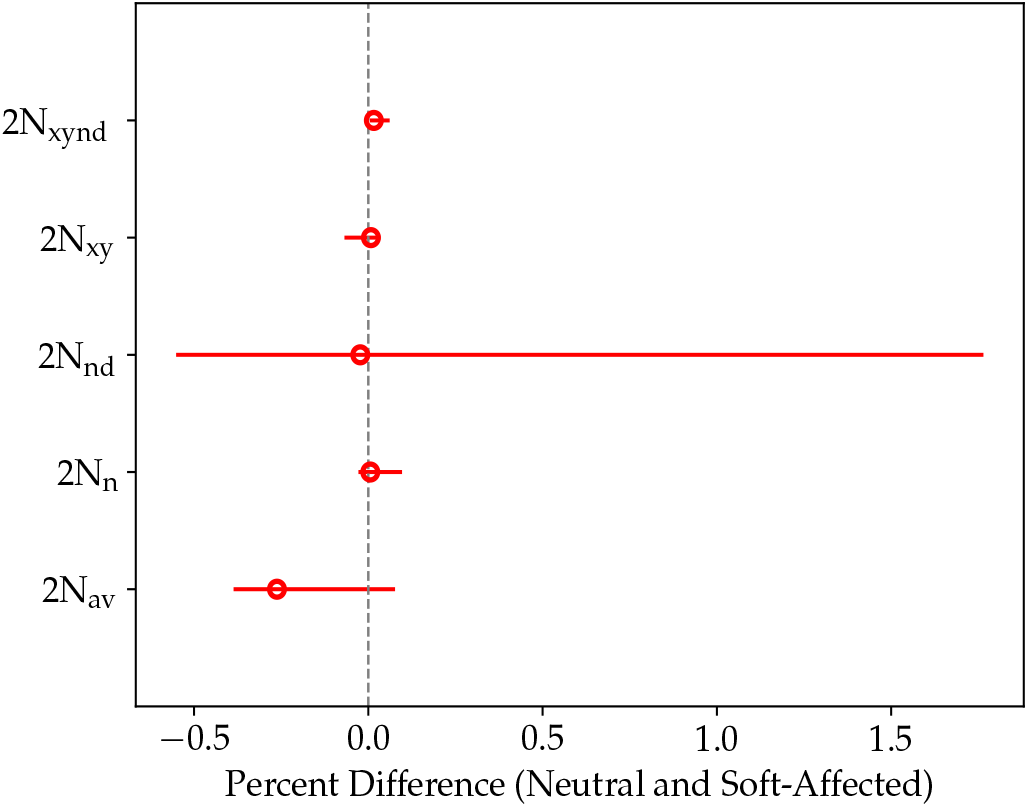
Percent difference between estimates of ancestral effective population size for soft-sweep associated regions and neutral regions for a model including Yorubans (*x*), European (CEU) (*y*), Neanderthals (*n*) further split into the Vindija (*v*) and Altai (*a*), Denisovans (*d*), and superarchaic hominins (*s*).

**Figure 4:**
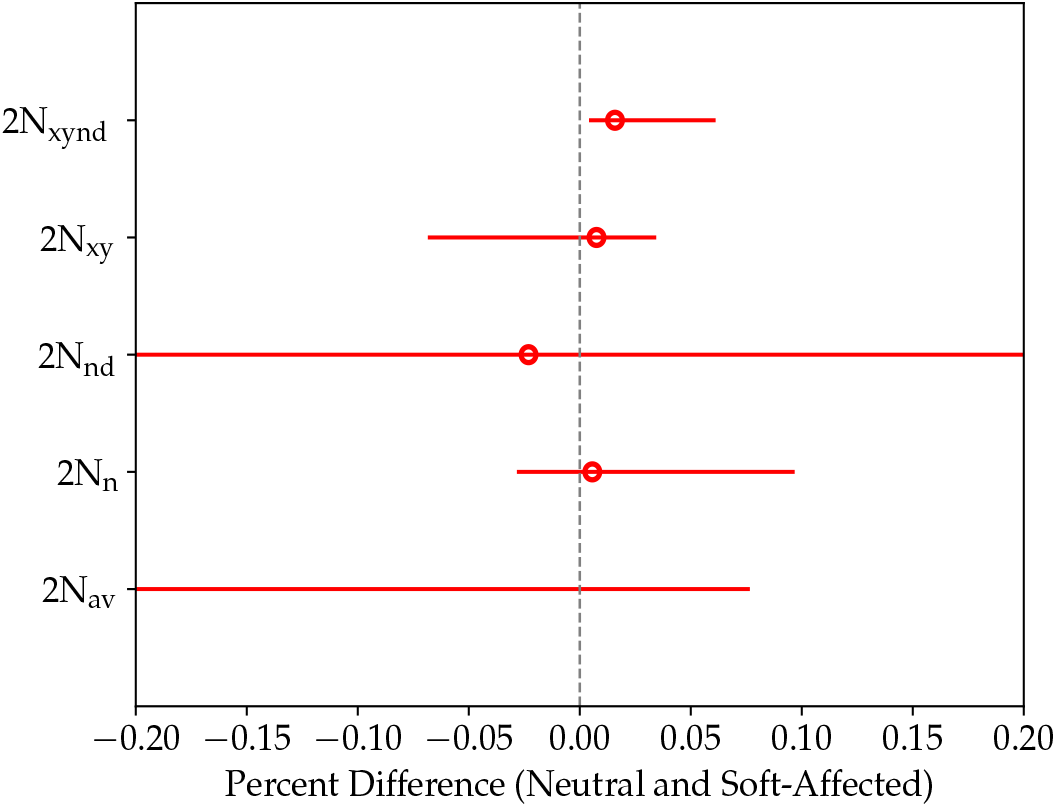
Percent difference shown in 3 zoomed in to compensate for the visual bias caused by the large amount of error around estimates of 2N_xynd_.

## 3 DISCUSSION

The difference in site pattern frequencies is in agreement with the results of Schrider and Kern (2017). The site patterns for the neutral regions imply a different population history than the selection affected regions. The neutral and selection-affected regions are close for each site pattern except the *x* and *xy* patterns. The *x* and *y* site patterns are relatively over-represented in the neutral case, while the *xy* site pattern is relatively underrepresented. The selection affected site patterns would therefore overestimate the length of the ancestral Eurasian branch, and underestimate the individual Eurasian population branches.

The significant differences in site pattern frequencies do not translate to large differences in estimations of population history in our model. In the European model, only one estimated parameter, the size of the ancestral population of humans, Neanderthals, and Denisovans (2N_xynd_), shows a significant difference between neutral and soft sweep affected regions. However, in this case the difference is more than an order of magnitude smaller than the estimated parameter itself.

The results here are consistent with large portions of the genome being under selection. However, differences in site pattern frequencies generated by soft selection do not appear to have substantial effects on the estimation of parameters of population history, at least when using *Legofit*. This result may have a larger effect at finer scales. For instance, the significant difference in T_av_, the divergence time of the Altai and Vindija Neanderthals, could be as large as 5,000 generations or 125,000 years. This may not too large when considering recent human population history and admixture between humans and other hominins, but could be problematic in reconstructing a fine scale history of Neanderthal populations. Further work exploring different time scales and population histories should be done before any generalization of these results is made. Nonetheless, soft sweeps do not cause meaningful disturbances in estimates of population history at the scale studied here.

## 4 METHODS

### Simons Data

Simons Genome Diversity Mallick et al. (2016) data for Japan (JPT), Yoruba (YRI), and Europe (England and France) was acquired from https://www.ebi.ac.uk/ena/data/view/PRJEB9586. Sites with a map quality or genotype quality below thirty were excluded. Sites that were fixed across all human populations were removed, because they cannot differentiate human populations. Indels, SNPs within seven bases of an indel, and low quality SNPs (filter level equal to zero) were removed. The Central Europeans from Utah (CEU) sample is not represented in the SGDP. Instead, individuals from England and France were used as a proxy for CEU. These samples were used because CEU is present in the analysis and results of Schrider and Kern (2017), but absent from the Simons data.

### Subdivisions

Schrider and Kern (2017) subdivide the genome into neutral and selection linked, soft-sweep affected, and hard-sweep affected regions. These results were obtained from https://github.com/kern-lab/shIC/tree/master/humanScanResults. The selection linked regions are those that are commonly assumed to be neutral in the literature, but showed evidence of being linked to and affected by selection in their machine learning analysis. The selection-affected and neutral regions were used here to divide data respectively. Regions inferred to be under selection are not studied here because selected regions are already expected to produce different estimates of population history from neutral regions. Here we are concerned with regions of the genome that could be mistaken for neutral regions and skew population histories in the literature. Each sample has its own subdivision that reflects the population and selective history of the population it belongs to. The neutral distinction made here refers to sites where all three populations are considered neutral. For a site to be included in the “selection-affected” for the purpose of this analysis, at least one population needs to be represented in the original “selection-affected” distinction.

### Site Pattern Frequencies

The program *sitepat* from the *Legofit* (Rogers, 2019) package was used to calculate relative site pattern frequencies (SPF) (Rogers et al., 2017). The ancestral allele is determined by using reference alleles of chimpanzees and gorillas. The analysis was limited to sites where the chimpanzee and gorilla were fixed for the same allele, and the human samples were polymorphic. One thousand bootstrap replicates were generated for each combination of selection type and set of populations. The difference in site pattern frequencies between selection affected regions and neutral regions was then taken, with confidence intervals generated from differences in individual bootstrap replicates.

### Legofit

Site pattern frequencies were used to estimate population history parameters using *Legofit*. The model of population history includes admixture between Eurasians and Neandethals, and between superarchaics and Denisovans and the ancestor of Denisovans and Neanderthals. Site pattern frequencies are generated using high coverage Neanderthal genomes from the Vindija (Prüfer et al., 2017) and Altai (Prüfer et al., 2014) Neanderthals, and a high coverage Denisovan genome from Siberia (Meyer et al., 2012). The Yoruban samples from the SGDP represent Africa. The final human population was a mixture of English and French individuals serving as a proxy for CEU. One thousand bootstrap replicates were generated for each model of population history. Confidence intervals for these differences were generated by taking the inner ninety-five percent of the differences between individual bootstrap replicate estimations. This process was conducted for neutral and soft-selection affected regions separately.

